# Higher Meta-cognitive Ability Predicts Less Reliance on Over Confident Habitual Learning System

**DOI:** 10.1101/650556

**Authors:** Sara Ershadmanesh, Mostafa Miandari, Abdol-hossein Vahabie, Majid Nili Ahmadabadi

**Affiliations:** School of Cognitive Science, Institute for Research in Fundamental Sciences, Tehran, Iran; Cognitive Systems Laboratory, School of Electrical and Computer Engineering, University of Tehran, Tehran, Iran

**Author notes:** These two authors contributed equally (Abdol-hossein Vahabie, Majid Nili Ahmadabadi).

## Abstract

Many studies on human and animals have provided evidence for the contribution of goal-directed and habitual valuation systems in learning and decision-making. These two systems can be modeled using model-based (MB) and model-free (MF) algorithms in Reinforcement Learning (RL) framework. Here, we study the link between the contribution of these two learning systems to behavior and meta-cognitive capabilities. Using computational modeling we showed that in a highly variable environment, where both learning strategies have chance level performances, model-free learning predicts higher confidence in decisions compared to model-based strategy. Our experimental results showed that the subjects’ meta-cognitive ability is negatively correlated with the contribution of model-free system to their behavior while having no correlation with the contribution of model-based system. Over-confidence of the model-free system justifies this counter-intuitive result. This is a new explanation for individual difference in learning style.

## Introduction

Learning is a crucial part of animals’ repertoire for survival. To address the wide range of problems that one encounters in the complex natural world, many animals, including humans, have developed multiple systems of learning and decision-making Daw et al. (2005); Dolan and Dayan (2013); Kahneman and Egan (2011). Different accounts of learning posit that there are at least two major systems that are contributing to the process of learning and decision-making. One system is habitual and reflexive, and the other is goal-directed and reflective Dolan and Dayan (2013). The reflexive system is computationally congenial, but it uses the experiences inefficiently, from a statistical standpoint. On the other hand, the reflective system uses experiences efficiently but requires substantially more computational resources Dolan and Dayan (2013). While this diversity in function enables animals to better handle the challenges which they face, it gives rise to the problem of arbitration between them. Different studies have shown that there is a considerable difference among people regarding their usage of these systems or as we call it, learning style. It seems that there is a connection between learning style and some cognitive capabilities.

One of the prominent ways to model this spectrum of learning style is Reinforcement learning theory Daw et al. (2011); Lee et al. (2014); Sutton et al. (2000). This theory states that the reflexive and reflective modes of behavior arise from two parallel value learning systems that are dissociable at the behavioral and neural levels. This RL model posits that the behavior is a combination of these two value systems. The crucial difference between these systems is whether or not they rely on an internal model of the environment. The reflective system, which is called model-based in RL framework, incorporates this model for evaluation of possible actions. On the other hand, the reflexive system, which is called model-free in RL framework, draws solely on the history of outcomes and seeks to repeat the rewarded actions.

Model-based and model-free methods, and their linear combinations, converge to the same policy after sufficient experiences in relatively stationary environments. Model-based method results in higher sample efficiency and faster learning in such environments, however, the advantages of the model-based method is not necessarily retained in non-stationary environments. Nevertheless, we witness variability in learning style in such environments where model-based and model-free methods have similar performances Kool et al. (2016); Akam et al. (2015). Here, the question is if any other property, except performance and learning speed, is potentially involved in the variability of the learning style. We were interested to see if these two learning methods differ in the second order decisions; i.e. confidence in decisions. If one of the methods results in a higher difference among decision values, it would induce higher confidence in decisionsAitchison et al. (2015); Folke et al. (2017).

Confidence is one of the commonly used paradigms to evaluate meta-cognitive ability Brainard and Vision (1997); Cheesman and Merikle (1986). Meta-cognitive ability is the ability to evaluate decisions Fleming and Daw (2017). This ability predicts higher (lower) levels of confidence in correct (wrong) decisions Maniscalco and Lau (2012). Therefore, we hypothesize that people with a higher level of meta-cognitive ability rely less on the learning mode that induces exaggerated confidence in decisions.

To examine this hypothesis, we use the two-step decision-making task developed by Daw et al.Daw et al. (2011), where both learning modes are the same in terms of performance due to non-stationary nature of the reward Daw et al. (2011); Akam et al. (2015). We also measure subjects’ meta-cognitive ability using retrospective confidence judgments in a word recognition memory test Sadeghi et al. (2017); Baird et al. (2013). We show that model-free strategy induces over-confidence in decisions, relative to model-based strategy, and meta-cognitive subjects rely less on the over-confident strategy.

## Results

Two tasks were used in this study. One was a word recognition memory task which was used to measure the meta-cognitive ability of subjects Baird et al. (2013); Sadeghi et al. (2017). The other was a sequential two-step decision-making task. It was designed so that the contribution of model-based and model-free components to the decision can be measured Daw et al. (2011). Forty-seven subjects completed the tasks. We paid subjects a fixed amount of money plus an extra reward proportional to their performances in both tasks. We investigated the relationship between behavioral characteristics in two tasks using statistical analysis and computational modeling.

### Two-step decision-making task

We adopted the two-step decision-making task from Daw et al. (2011)Daw et al. (2011) (Figure 1). The task was tailored around a treasure hunt story. In the story, the subjects had a choice between two alternative airplanes that would take them to either of two jungles where they could hunt for gold. Subjects completed 200 trials of this task. At each trial, they selected between two options that led probabilistically to either of the two second-step states. Each option of the first step was linked to one of the second-step states and led there 70% of the times. After that, the subjects chose between two alternatives in the second-step state, followed by a probabilistic binary reward. To ensure continuous learning through the task, the reward probabilities were non-stationary; changed steadily and independently from each other during the task.

**Figure 1:**
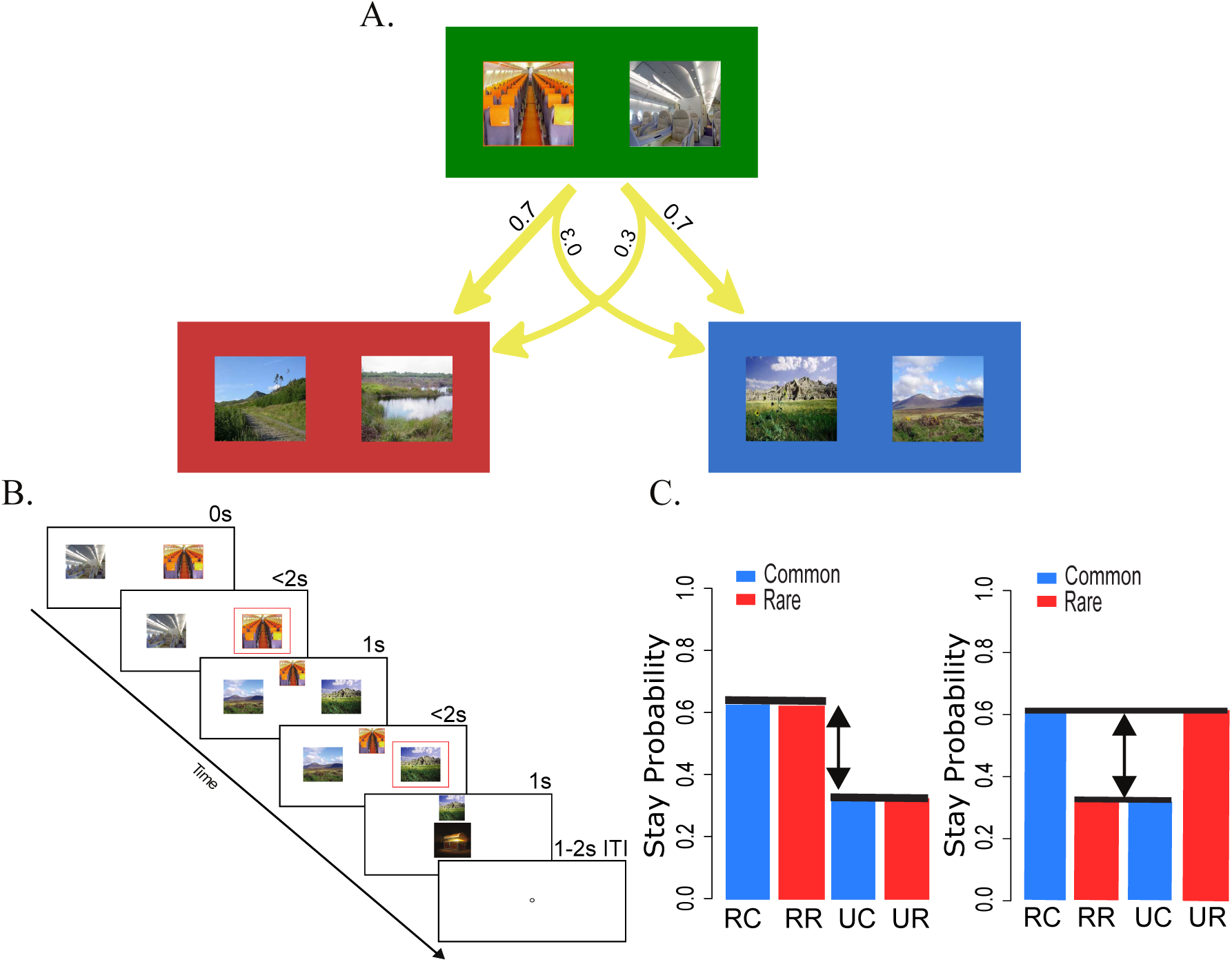
Two-step decision-making task. A) Each first-step action was commonly associated with the transition to one of the second-step states. Although, it also could lead to the other second-step state, but rarely. B) The timing of a single trial: a choice in the first step followed by a choice in the second step. The second choice was reinforced by the reward. C) Left, model-free RL predicts a high probability of repeating (stay probability) the first-step choice of the previous trial if it is rewarded. Type of transition, common or rare, is not effective on the stay probability in this case. Right, model-based RL predicts that the type of transition is effective on the stay probability. Thus model-based learning is influenced by the interaction of reward and transition.

We studied the effect of reward (model-free component), also the interaction of reward and transition (model-based component) in the subjects behavior in the two-step task. To indicate the model-based and model-free components, we studied the effects of two main factors, outcome (rewarded or unrewarded) and transition type (common or rare) of each trial, on repeating the first-step choice in the next trial (we call probability of this event P-stay). As it was depicted in Figure 1, the reward is the only predictor of P-stay of a pure model-free agent while for a pure model-based agent it is only the interaction of reward and transition that contributes in the prediction of P-stay. Applying the repeated-measures ANOVA indicated that a combination of the two strategies, model-based and model-free, was involved in guiding the behavior (main effect of reward: *F* (1, 36) = 23.5, *p* = 2 × 10^−5^, interaction between reward and transition: *F* (1, 36) = 13.11, *p* = 0.0008). As in Daw et al. (2011)Daw et al. (2011), we concluded that both reward and task structure are employed by the subjects.

### Memory task

The memory task was a Persian word recognition memory test based on Baird et al. (2013)Baird et al. (2013) and adapted from Sadeghi et al. (2017)Sadeghi et al. (2017). The task had two phases: encoding and recall. During the encoding phase, 140 words out of a set containing 280 words, were presented on the screen, separately and sequentially. In the recall phase, immediately after the first phase, all 280 words were displayed sequentially. subjects decided if each word had been seen in the encoding phase or not. After that, they reported their confidence about the correctness of their response on a scale of 1 (low confidence) to 6 (high confidence)(Figure 2). Individuals did not receive any feedback about the correctness of their response.

**Figure 2:**
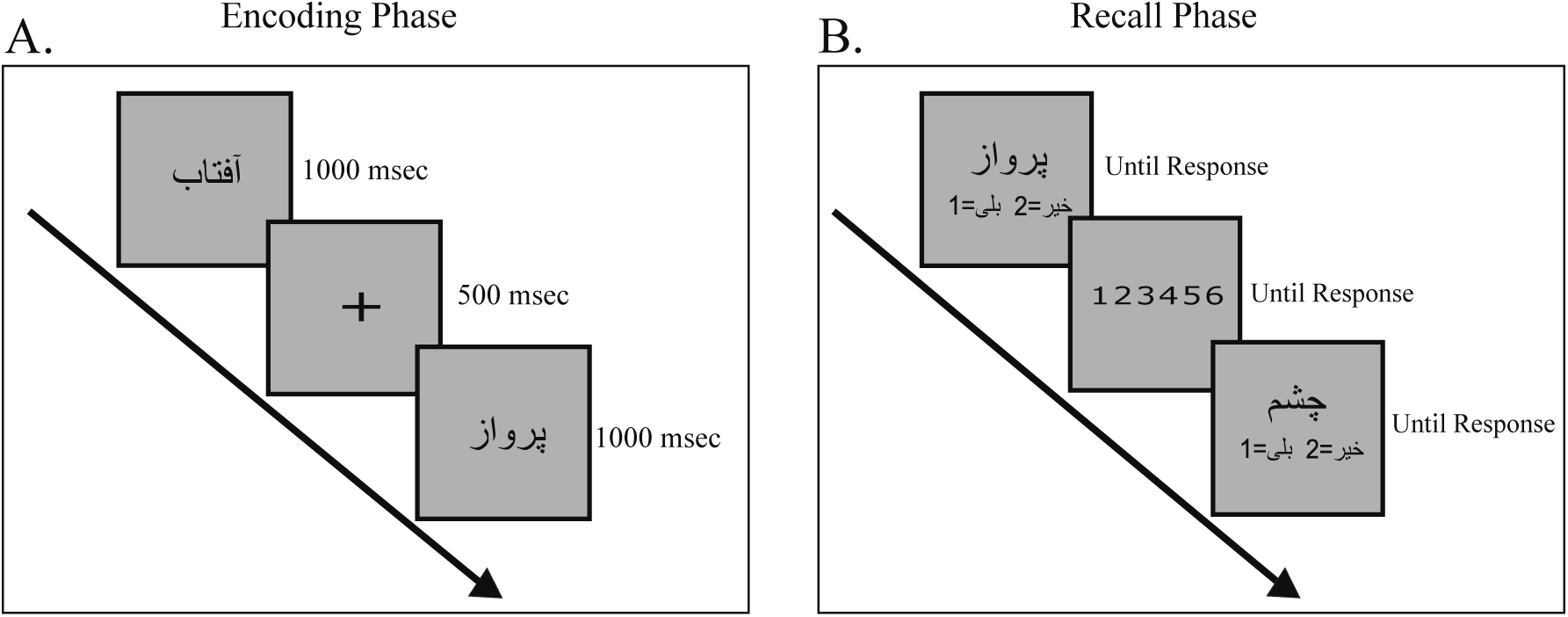
Memory task. This task had two phases: 1) In the encoding phase, we asked the subjects to memorize half of the words which were chosen randomly from a word set. 2) In the recall phase, after the presentation of each word from the word set, the subjects reported whether the word was seen in the encoding phase or not. After the response, they scaled their confidence about the previous response.

In the memory task, average memory performance of our subjects was 70% (SD 0.07, range 0.57-0.86), comparable to the results reported in Baird et al. (2013) Baird et al. (2013), 71% (SD 0.09, range 0.57-0.91). Reported results about meta-cognitive accuracy were measured by M.Ratio method with unequal-variance assumption Maniscalco and Lau (2012). However, we had the same results by both M.Ratio and HMeta-d methodsFleming (2017) which both have an equal-variance assumption.

## The impact of meta-cognitive ability on learning style

To study whether the meta-cognitive accuracy predicts learning style, we measured these two characteristics in the memory and the two-step decision-making tasks accordingly.

We investigated the relationship between model-based/model-free components and meta-cognitive accuracy using two methods of analysis. Mathematical details of these methods are described in the methods section. The results from these two methods are described below.

### P-stay analysis

We measured model-based and model-free components of behavior through measuring the probability of first-step decisions in two consecutive trials. There are four possible conditions in every trial; a multiplication of 2×2 conditions: common/rare transition and reward/no-reward conditions. The four conditions are named rewarded common (RC), rewarded rare (RR), unrewarded common (UC), and unrewarded rare (UR). The P-stay plot for pure model-based and model-free agents are shown in Figure 1. Model-based and model-free indices were calculated by Eq1 and Eq2 Miller et al. (2016); Eppinger et al. (2013):

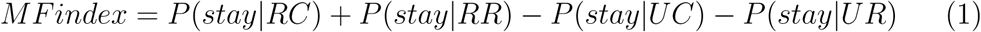

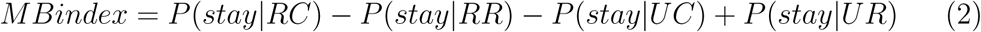

The model-free index is the difference of stay probabilities between rewarded and unrewarded conditions and indicates the effect of reward (model-free component) on the behavior. The model-based index is the difference of stay probabilities between two conditions; the first condition consists of RC and UR, and the second condition consists of RR and UC.

In pure model-free learning, task structure is not employed, and the reward is the predictor of P-stay. Therefore, the difference of P-stay in rewarded and unrewarded conditions is the *MFindex*. In contrast, in pure model-based learning, both reward and task structure are used, so reward × transition is the predictor of P-stay. Hence the difference of P-stay in two conditions, related to two quantities of reward × transition is the *MBindex*. Two-sided arrows in Figure1 (C and D) visualize how *MFindex* and *MBindex*, is calculated respectively.

**Figure 3:**
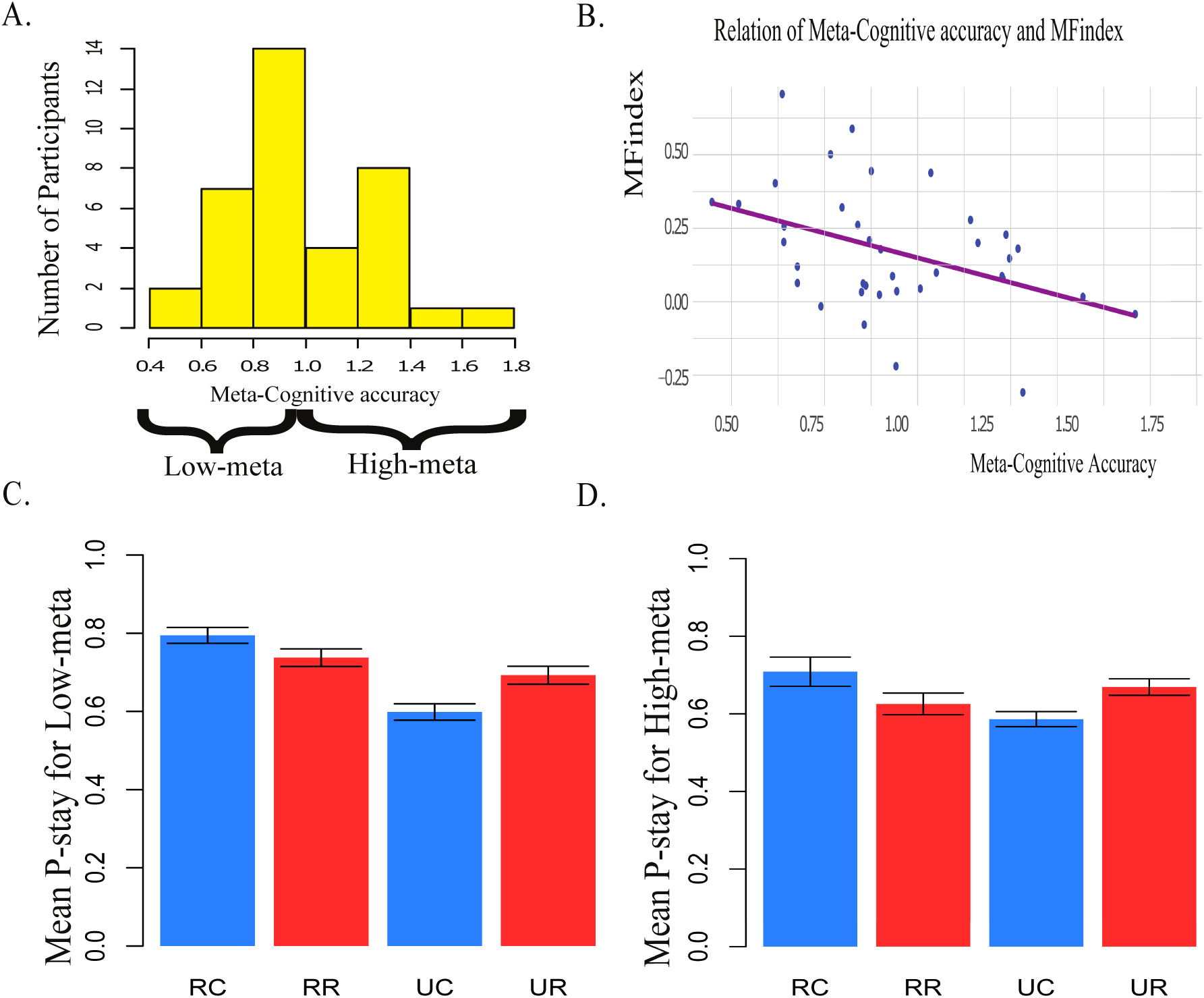
(A) The histogram displays meta-cognitive accuracy in Memory task’s data. Samples are divided by a median split into two groups. (B) Scatter plot shows meta-cognitive accuracy (x-axis) versus *MFindex* in P-stay analysis (y-axis). The slope of the fitted line shows the correlation of model-free behavior and meta-cognitive ability. (C)(D) The P-stay for Low meta-cognitive and high meta-cognitive groups, as you can see, there is a clear difference between the two. We affirmed the significance of this difference by statistical measures on *MFindex* and *MBindex*.

*MFindex, MBindex*, age and memory performance were used as the regressors to predict meta-cognitive accuracy. We obtained significant negative coefficient for *MFindex* (−0.635, *p* = 0.003) while the role of *MBindex* was not siginificant (0.068, *p* = 0.661). Age did not have any significant role in prediction of meta-cognitive accuracy (−0.003, *p* = 0.762) and the memory performance had a significant negative coefficient (−1.608, *p* = 0.004). We did not interpret negative coefficient of performance as a regressor, because this result did not remain with different measures of meta-cognitive accuracy.

### Modeling

For a more in-depth analysis of behavior, we employed computational modeling. While P-stay analysis has the advantage of being relatively model agnostic Daw et al. (2011), behavior in each trial is analyzed only by looking at its previous trial, so it is beneficial to have a more encompassing analysis that incorporates more of the history in analyzing behavior. For this, we used reinforcement learning modeling similar to previous studies Daw et al. (2005, 2011); Kool et al. (2016). In this model, the model-free and model-based components update the value of each action in parallel, and the estimated values then will be linearly combined (Eq5). The combined values then go through a softwax function that calculates the probability of choosing each action. For estimating the value of each action, the model-free system relies only on previous rewards while the model-based system uses both the rewards and transition probabilities for value estimation. Free parameters of the model are: learning rate *α*, softmax temperature *β*, perseverance *p*, eligibility trace *λ*, direction bias *s*, and the proportion of model-based to model-free control, *ω*. We used correlation test between meta-cognitive accuracy and *ω* (Pearson correlation = 0.474, *p* = 0.002, 95%*CI* = [0.178, 0.692]). It means that meta-cognitive ability predicts the relative proportion of model-based and model-free components in decision-making.

However, it was unknown whether the meta-cognitive ability is correlated with either model-based or model-free strategies, or to both. We cannot answer this question by the models presented in previous studies Daw et al. (2011); Akam et al. (2015); Kool et al. (2016) because in these models, a weighted sum of model-based and model-free components was used for decision-making. Therefore, we removed the restricting weighted sum condition and applied two independent weights (*ω*_*mb*_ and *ω*_*mb*_) for the linear combination of model-based and model-free values; see Eq7 in the methods section. Also, this model was inspired by neural evidence from Lee et al. (2014)Lee et al. (2014), so we hypothesized that it will better capture the subject’s behaviors. *ω*_*mb*_, *ω*_*mf*_, the age of the subjects and their performance in the memory task were used as regressors to predict meta-cognitive accuracy. We obtained significant negative coefficient for *ω*_*mf*_ (−0.530, *p* = 0.001) while the role of *ω*_*mb*_ was not significant (0.013, *p* = 0.909) (Figure 4, B and C). Age did not have any significant role in prediction of meta-cognitive accuracy (−0.003, *p* = 0.746) and memory performance had a significant negative coefficient (−1.678, *p* = 0.003). Similar to the P-stay analysis, we did not interpret the negative coefficient of performance as a regressor, because this result did not remain with different measures of meta-cognitive accuracy like HMeta-d. Moreover, as we hypothesize, this model fitted to the subject’ s behavior better than the original model from Daw et al. (2011)Daw et al. (2011). The average BIC was 4.6 units lower for our model.

We were interested to know what is the difference between model-based and model-free strategies that can be a candidate explanation for why the high meta-cognitive subjects rely less on model-free strategy. Thus we computationally studied these two strategies in the two-step decision-making task.

**Figure 4:**
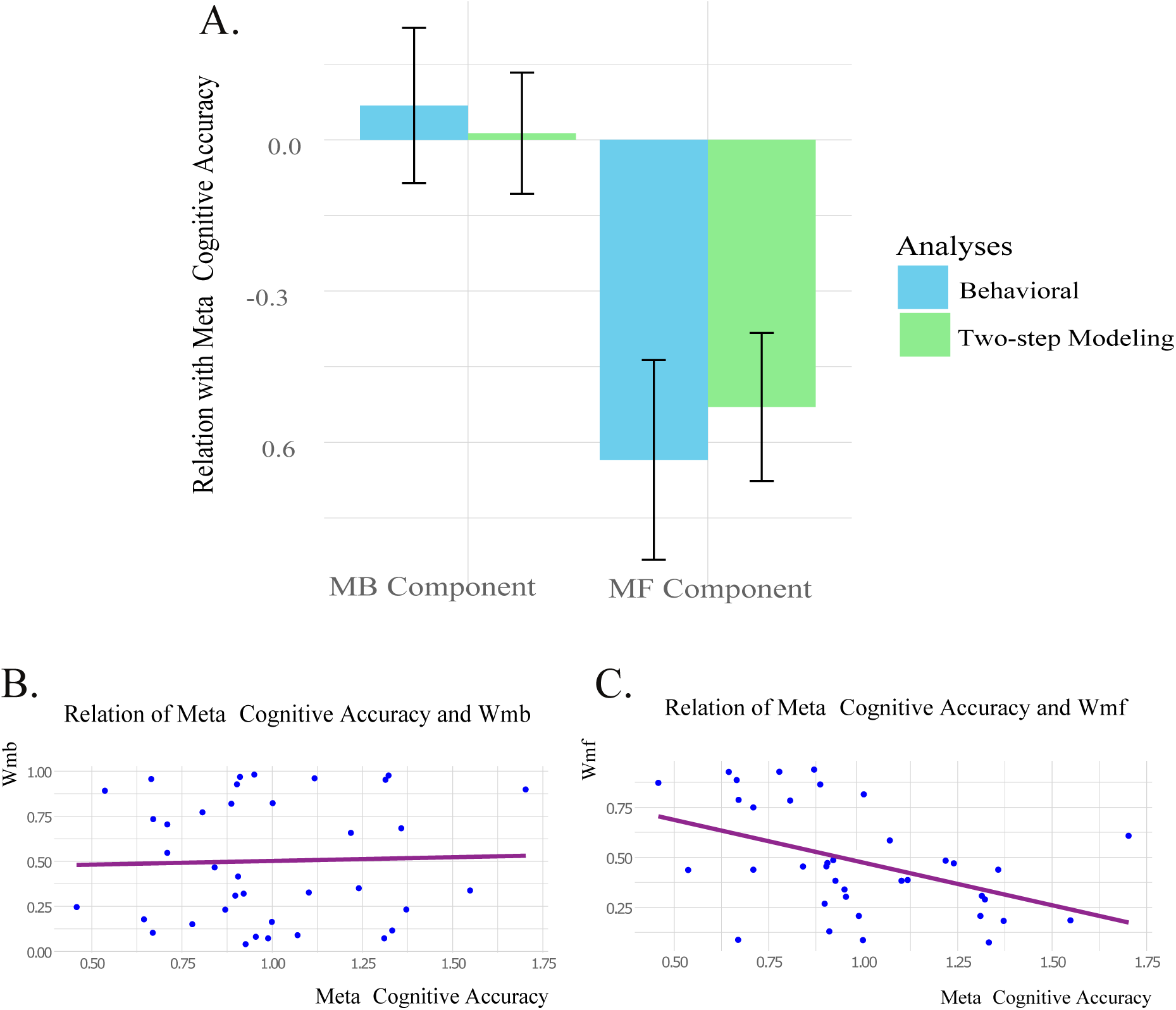
A) The meta-cognitive accuracy is predicted with model-based and model-free components in two analyses. Two regression analyses were used, in each of them regressors were from one of the two analyses; P-stay analysis (blue) and reinforcement modeling (green). B) Scatter plot shows meta-cognitive accuracy (x-axis) versus model-based component in the modeling analysis (y-axis). The slope of the fitted line shows the correlation of the model-based behavior and the meta-cognitive ability. C) Scatter plot shows meta-cognitive accuracy (x-axis) versus the model-free component in modeling analysis (y-axis). The slope of the fitted line shows the correlation of model-free behavior and the meta-cognitive ability.

## Learning style and performance analysis

Similar to the previous studiesAkam et al. (2015); Kool et al. (2016), our subjects’ performance in the two-step task was not significantly better than the chance level (Mean = 0.473, t(36) = −4.106, p = 0.999, 95%*CI* = [0.473, 0.500]). According to Akam et al. (2015) Akam et al. (2015), neither model-based nor model-free agents cannot act better than a random agent in this task setting. Moreover, both model-based and model-free indices were not significantly correlated with the performance in the two-step task (Pearson correlation = 0.259, *p* = 0.1202, 95%*CI* = [−0.069, 0.538] and (Pearson correlation = 0.135, *p* = 0.423, 95%*CI* = [−0.196, 0.440] respectively. As a result, something beyond performance should be involved in the learning style. Here, we investigate possible role of second order decision.

### Confidence correlate of model-based and model-free strategies

According to Folke et al. (2016)Folke et al. (2017), confidence can be strongly predicted by the difference of action values; the same is reported in perceptual decision-making where the difference in evidence is correlated with confidence Aitchison et al. (2015). We exploited the same concept and checked if model-based and model-free methods differ in estimating the difference of Q-values for first-step choices. To see the overall trend, we used reinforcement learning model fitting to estimate the Q-values of first-step actions for each subject Daw et al. (2011); Kool et al. (2016). To compare confidence correlate of model-based and model-free strategies, we measured the absolute difference of first-step Q-values for each strategy in every trial. For hypothetically pure model-based and model-free subjects we have:

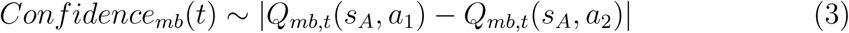

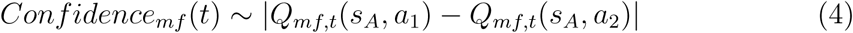

*Q*_*mb,t*_(*s*_*A*_, *a*_1_) is Q-value of model-based strategy in trial *t* for one of the options in the first-step and *Q*_*mb,t*_(*s*_*A*_, *a*_2_) is for the other one. The same is for corresponding Q-values for model-free strategy.

Confidence for the model-free strategy (Mean = 0.2528) was significantly higher relative to model-based strategy (Mean = 0.1214, t(45.956) = −6.2952, p < 1.048*e -*07). Considering the fact that neither model-based nor model-free methods act better than a random agent, relying more on the model-free strategy results in lower meta-cognition due to the exaggerated confidence it induces. It is noticeable that variances in the Q-values of each first-step decisions were not significantly different for model-based and model-free strategies (*F* (36) = 1.314, *p* = 0.415, 95%*CI* = [0.677, 2.553] and *F* (36) = 1.450, *p* = 0.269, 95%*CI* = [0.746, 2.816] respectively). Thus, the negative correlation of meta-cognition and model-free index cannot be explained by the variance of Q-values.

We also showed with mathematical calculations that the difference in Q-values is bigger for the model-free relative to the model-based method in every trial on the base of our fitted parameters; see supplementary materials.

**Figure 5:**
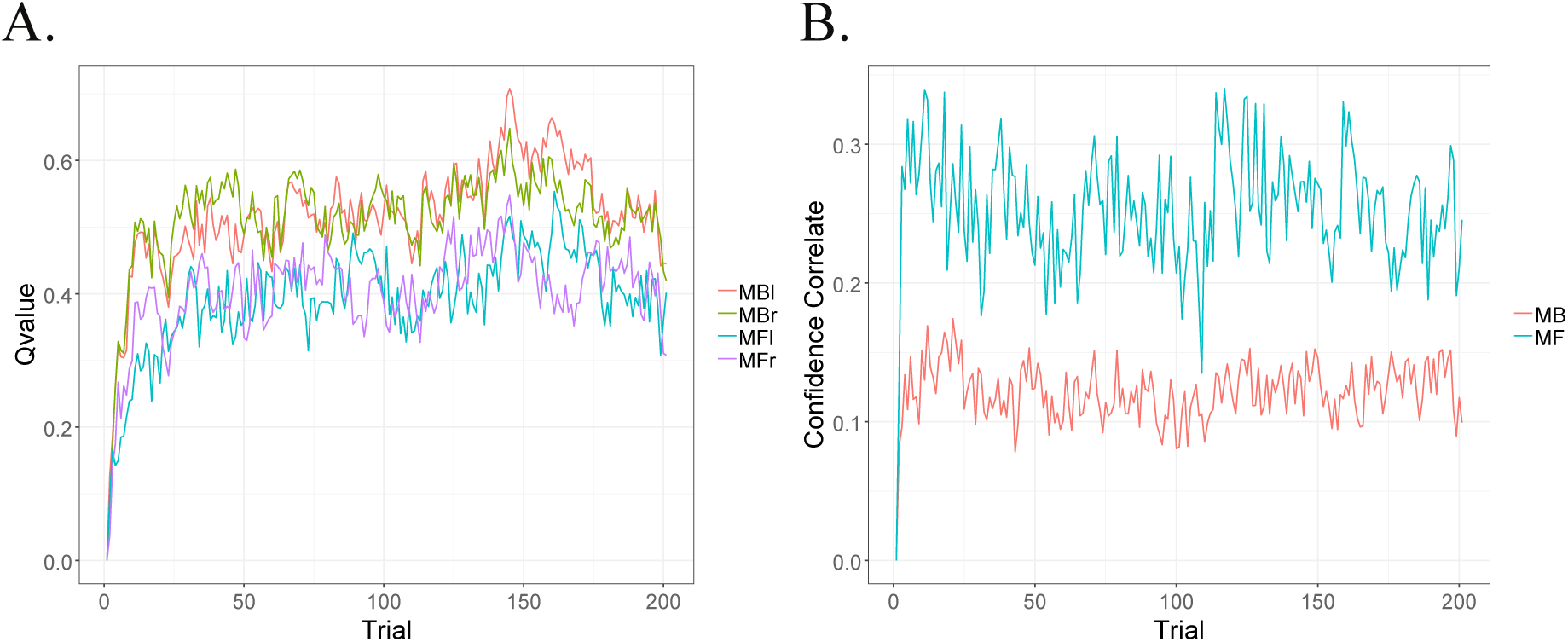
A) The estimated Q-values of two options in the first-step over trials for model-based and model-free strategies. The difference of Q-values are higher in the model-free strategy. B) Corresponding confidence of the model-based and the model-free modules measured by absolute difference between the Q-values of two options. Confidence correlate was significantly higher in the model-free strategy.

**Figure 6:**
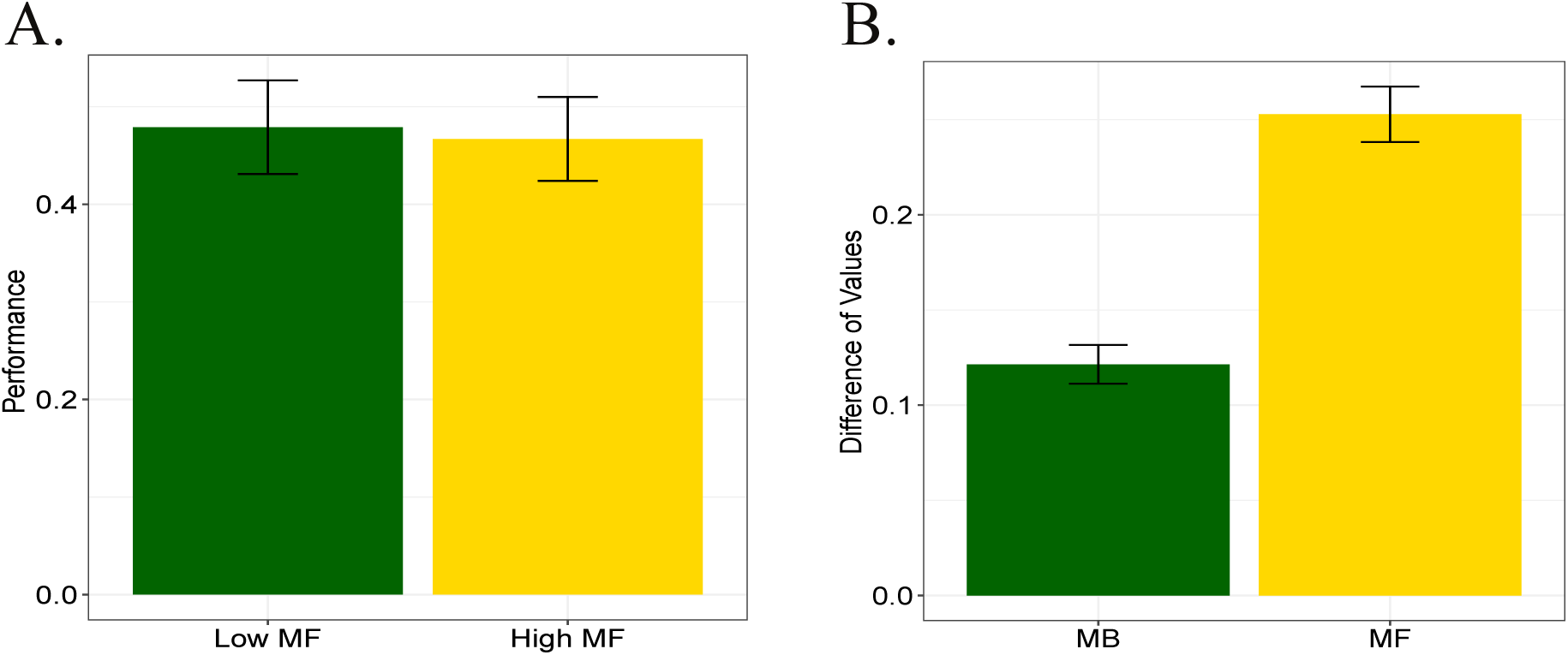
A) Performance of the subjects in the two-step decision-making task was not significantly correlated with their model-free index; i.e. neither the high model-free subjects nor the low model-free ones had significant different performance. B) The mean confidence correlate of subjects during trials if they behaved according to the model-free strategy was significantly higher relative to their mean confidence if they fully relied on the model-based strategy.

## Discussion

In the two-step decision-making task setting which we used in this paper, model-based and model-free learning strategies as well as a random decision maker had the same performance and learning speed. However, we observed the individual difference in the subjects learning style; the subjects were different in their model-free and model-based indices. Our question was if any other property, far from performance and learning speed, is associated with this individual difference. We were interested to see if these two learning strategies are different in the second order decisions, i.e. confidence in decisions. According to Folke et al. (2016) Folke et al. (2017), the difference in action values is a confidence correlate. We showed that the model-free system had exaggerated confidence correlate in decisions while having no difference in performance with the model-based method; i.e. higher reliance on model-free learning results in lower meta-cognition. In line with this analysis, we observed that there is a strong negative correlation between model-free behavior of our subjects and their meta-cognition level in a word recognition memory task while having no correlation with model-based behavior. Lee et al. Lee et al. (2014) showed that the degree of model-based control on decision-making could negatively modulate the connectivity between the arbitrator and model-free valuation regions of the brain. We suggest that there are some similarities between this finding and the negative correlation we observed in our results. The subjects with high meta-cognitive ability may have relied on the more meta-cognitive system, the model-based one, by attenuating the contribution of the less meta-cognitive system; i.e. the model-free system.

The commonly used RL modelDaw et al. (2011) does not explain our behavioral data, because the model assumes sum-to-one weights for model-based and model-free Q-values in a linear combination. According to that model, the strong negative correlation between meta-cognition and model-free index should accompany a positive correlation of meta-cognition and model-based index; which was in contrary with our behavioral data. Therefore, we assumed independent weights for model-based and model-free modules. The model-fitting result was in-line with our data and the BIC was lower even in the face of having more parameters in comparison with the original model. We also had biological inspiration for this model. Lee et al. (2014) Lee et al. (2014) reported a significant negative correlation between the activity of arbitration region and the strength of the connection between putamen and vmPFC. The putamen is a candidate region for model-free behavior and vmPFC is a candidate region for combining Q-values of the two learning systems. In contrast, they did not report any significant correlation between the activities of the arbitrator regions and the strength of the connection between candidate regions for model-based behavior and vmPFC. This result suggests that there is a difference between the relation of arbitration regions activity and connectivity of model-based and model-free regions with the Q-value integration brain part. Combining our modeling results and the neural evidence from Lee et al. (2014) Lee et al. (2014), we assert that our model with independent weights can better depict the human learning behavior and hence can supersede the previous model in the future works. Otto et al. (2014) showed that cognitive control is correlated with model-based behavior and not with model-free one. This result may seem contradictory to our result, but we argue that they can co-exist. As Shea et al. (2014) Shea et al. (2014) discussed, one could have high cognitive control but low meta-cognitive ability, and meta-cognitive ability is not essential for every type of cognitive control, especially cognitive control which requires attention to cues and quick responses. The cognitive control measured in Otto et al. (2014)Otto et al. (2014) was of this kind; this type of cognitive control does not involve the kind of decision evaluation that is required for assessing decisions in our memory task. Hence, the apparent difference between our results and those of Otto et al. (2014)Otto et al. (2014) may be rooted in possible differences of meta-cognitive accuracy and cognitive control Shea et al. (2014). Nevertheless, further studies are needed to clarify the details of meta-cognition and cognitive control relation with learning style.

It is important to emphasize that the correlation we found, does not imply any direction of causality. In addition, we cannot reject the possibility of a third cognitive ability that influences both meta-cognitive ability and learning style. Further studies are needed to establish a causal relationship. If meta-cognition is the higher node in the hierarchy of control in the decision-making process, it might be the case that neuromodulation manipulation of meta-cognition Hauser et al. (2017) affects the learning style. It has been also shown that learning style can be influenced by neuromodulator administration Wunderlich et al. (2012). Measuring the change of meta-cognition in learning style manipulation experiments will help to get a clearer picture.

The low meta-cognitive ability has been reported in some psychiatric disorders, while the memory performance of subjects was intact Rouault et al. (2018). Considering our finding about the relation of meta-cognitive ability and model-free behavior, it is possible to investigate whether people with the mentioned disorders would exhibit a bigger reliance on model-free behavior than normal. The same can be studied about psychological disorders that are correlated with an excess of model-free influence on decision-making process Dolan and Dayan (2013); Gillan et al. (2011); Maia et al. (2008). It can be studied whether people with these disorders show any sign of impairment in their meta-cognitive ability. If future studies confirm the existence of mentioned relationships, they could play a role in developing new treatments for these disorders.

## Methods

### Subjects

47 subjects (23 females, aged 19-32, mean age = 23.91) completed the tasks. According to previous studies Decker et al. (2016); Eppinger et al. (2013) that found a large effect (*η*^2^ = 0.2) of individual differences in learning style, it was expected that the sample size of 36 would be enough to achieve 90% power to detect a true effect of a comparable size, if we assume *α* of .05. The subjects gave their informed consent, and the study procedures were approved by the ethical committee of the University of Tehran. We paid subjects a fixed amount of money plus an extra reward proportional to their performances in both tasks.

### Excluded subjects

Nine subjects excluded because their behavior was in conflict with an intention to acquire reward Decker et al. (2016); Otto et al. (2013). We had two criteria to check for this intention; the first criterion was if subjects showed a low tendency (less than 55% of the times) to choose the rewarded option from the previous trial in the case that they reached the same second step. The second criterion was if their behavior was best described by a random agent (an agent that make a decision with an equal probability between the available choices, regardless of anything) rather than a reinforcement learning agent. One subject only chose a single rate for her confidence more than 95% of trials in the memory task, so she was excluded Folke et al. (2017). Thus thirty-seven subjects remained for analysis. **Tasks and procedures.** Both tasks were computerized and programmed in MATLAB (Mathworks). For the memory task, COGENT 2000 toolbox was used (http://www.vislab.ucl.ac.uk/cogent.php). The two-step decision-making task was implemented with Psychophysics toolbox extensions Kleiner et al. (2007); Brainard and Vision (1997); Pelli and Vision (1997). **Memory task.** The task consisted of two encoding and recall phases. subjects saw 140 words one by one. Words were randomly selected for each subject from the word set. Each word was presented in the middle of the screen for 1000 ms, with 500 hundred inter-stimuli interval (ISI). During the ISI, a fixation point was shown in the middle of the screen. After the encoding phase, 280 words presented in the recall phase which half of them were the words in the encoding phase. subjects were instructed to report whether each word was in the encoding phase or not (old or new) and pressed key numbers 1 or 2 on the top left side of the keyboard accordingly. The subjects’confidence about the correctness of their decision were reported through numbers 1 (low confidence) to 6 (high confidence) on the keyboard (Figure 1). The experiment did not include any feedback to subjects about their responses. There was no time limit to answer either the recall questions or the confidence rating. **Two-step decision-making task.** The task was originally based on the task applied in Daw et al. (2011)Daw et al. (2011) (Figure 2). All subjects read a detailed description of the task. The task was about finding treasures by traveling to different places. The subjects chose in every trial between two airplanes that took them to a couple of different jungles where they could find treasures. The story was briefed for them by a Powerpoint presentation containing an animation version of the task. Subjects also answered a number of questions to make sure they could recognize and discriminate the pictures in the task from relatively similar pictures. They also completed 40 training trials before the start of the main task. The task consisted of 201 trials. At each trial, there were two left-right randomized options that led with different probabilities to one of the two second-step states. Each option of the first step was associated with one of the second-step states 70 % of the times. In second step of each trial, the subjects chose between two other options which led to a probabilistic binary reward. To keep continuous learning about the best option, reward probabilities were changed in each trial independent from each other. Four different Gaussian random walks were applied to change the reward probabilities of options. Response time in each step was limited to 2secs. Inter-stimuli intervals were 1 to 2secs, and between two trials there was 1sec meanwhile. Feedback was presented for 500 msecs.

## Data Analyses

### Meta-cognitive accuracy

Meta-cognitive accuracy was measured for each subject through meta-cognitive ratio (M.Ratio) with unequal-variance assumption Maniscalco and Lau (2012). This method was implemented in MATLAB using the toolbox provided by Maniscalco et al. (2012) Maniscalco and Lau (2012). M.Ratio is based on a Signal Detection Theory model Baird et al. (2013); Macmillan and Creelman (2004). The main important point in the measurement is that choices and confidence reports are inputs for two different SDT models, type 1 and type 2 respectively. Thus two different parameters *d′* and *metad′* are used to measure sensitivity in choices and confidence rating respectively. *d′* measures the accuracy of choice and *metad′* is a quantitative representation for the accuracy of the choices’evaluation. Meta-cognitive accuracy is the ratio of *metad′* to *d′*, measuring the amount of information used at the meta-level relative to the object level. In the memory task, the two kinds of stimuli (old and new, described in the Method section), are not necessarily modeled by equal variance SDT, so we reported our result with unequal variance SDT model of *metad′* Maniscalco and Lau (2012). We also checked our results with M.Ratio which has an equal-variance assumption. Furthermore, we repeated the analyses through hierarchical Bayesian estimation of meta-cognitive efficiency, known as HMeta-d method Fleming (2017). The Fleming’s method provides opportunities to enhance statistical power Fleming (2017). This method is based on M.Ratio with equal-variance assumption.

### Contribution of model-based and model-free components to decision-making

To replicate previous studies, we looked at whether there is evidence for the contribution of both systems to decision-making or not. For this purpose, the probability of repeating the first-step choice of the previous trial (P-stay), conditional on the transition type (common or rare) and feedback of the previous trial (reward or no reward), were entered in a two-way repeated-measures ANOVA Wunderlich et al. (2012). We studied the effect of reward (model-free component), also the interaction of reward and transition (model-based component) in the behavior. The analysis was performed in R programming language.

### Computational modeling based on reinforcement learning

To measure the contribution of model-based and model-free learning in the behavior, as in previous studiesDaw et al. (2011); Kool et al. (2016); Wunderlich et al. (2012), we used a hybrid parametrized model. In this model, model-based and model-free systems update their estimation of each action values on a trial-by-trial basis Daw et al. (2011); Kool et al. (2016). Then a weighted sum of these two values (Eq5) goes through a softmax function to form decisions (Eq6). The weight (*ω*) represents model-based component’s contribution to value-based decision-making. This weight is the proportion of model-based component to model-free component in the final value for choosing the first-step actions.

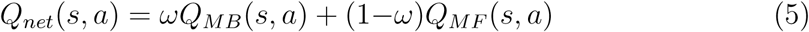

To connect *Q*-values to the probability of choosing one first-step action, softmax rule is used:

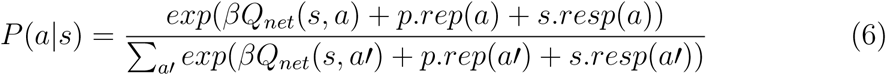

*β* is the inverse temperature, representing randomness in decision-making. Lower *β* leads to more uniform probabilities for different actions. *p* represents the degree to which subjects show perseveration (*p* > 0) or switching (*p* < 0) at the first step. The variable *rep*(*a*) is 1 if *a* is a first-step action and is the same as the action on the previous trial, and is 0 otherwise. The variable *resp*(*a*) is 1 if *a* is a first-step action which is selected by a response key the same as the key that was pressed on the previous trial and is 0 otherwise. *s* represents the degree to which subjects repeat (*s* > 0) or change (*s* < 0) the key presses at the first step of each trial.

This model has 6 free parameters (*β, α, λ, p, s, ω*), in which *λ* is an eligibility trace Sutton et al. (2000). Eligibility trace is employed to use the reward in the second step to update the values of actions in the first step.

Beside the mentioned model (Eq5), we developed a new model (Eq7) which draws on available neural evidence by Lee et alLee et al. (2014). They showed that the arbitration system incorporates the action values of the model-based and the model-free systems differently. Inspired by this result we hypothesize that the weights are determined independently from each other. Hence, we decided to relax the weighted average condition of the previous model and have independent weights for each system. We hypothesize that this model will fit the behaviors of our subjects better and also enable us to dissociate between the roles of model-based and model-free systems.

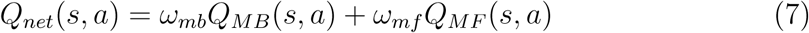

### Relationship between meta-cognitive accuracy and model-based/model-free components

To study the possible relationship between the meta-cognition and learning style, the Pearson correlation was performed between *w* and meta-cognitive accuracy. We also applied regression analysis to study whether model-based or model-free components predicts meta-cognitive ability. The regressors were *ω*_*mf*_, *ω*_*mb*_, the age of the subjects, and their performance in the memory task. The aforementioned analysis was repeated with *MBindex* and *MFindex* as regressors instead of *ω*_*mf*_, *ω*_*mb*_.

## Supplementary

### Experience effect on model-based and model-free learning

We investigate the effect of a single trial experience on the difference of first-step action values. Assume that at trial *t* the agent has chosen *a*_*t*,1_ in the first-step (*s*_1_), reached *s*_*x,t*_ (either *s*_2_ or *s*_3_), then chooses *a*_*t*,2_ and receives reward *r*_*t*_. Here, we calculate the difference in value update (Δ*Q*) of the first-step actions after this trial. The diagram of this task is shown in Figure S1.

After each trial the model-free updates its first-step action value according the formula below:

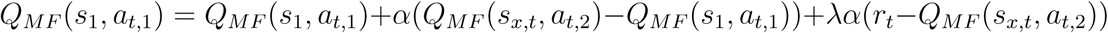

Where *α* is the learning rate, *λ* is the eligibility trace and Q(s,a) is the current value of action a in state s. Also, it is important to emphasis that only the executed first-step action value is updated after each trial. For the model-based agent we have:

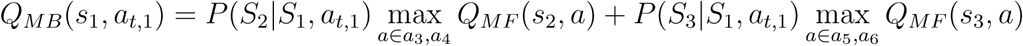

Considering these formulas, we can now compute the difference of first-step action values caused by each trial. For the model-free agent we have:

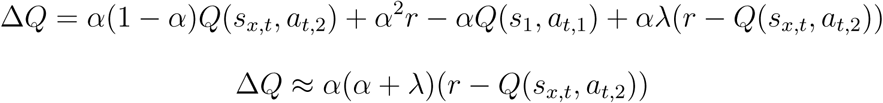

For the model-based agent in our task we have:

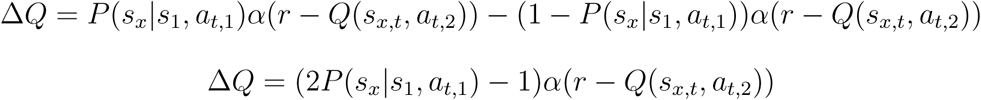

Where *P* (*s*_*x*_|*s*_1_, *a*_*t*,1_) is 0.7 for the common transition which makes the (2*P* (*s*_*x*_|*s*_1_, *a*_*t*,1_)− 1) equal to 0.4. For our task where the discount factor is one, the reward for transition from the first-step to the second step is zero and the difference in estimated action values is very small, *Q*(*s*_*x,t*_, *a*_*t*,2_) *- Q*(*s*_1_, *a*_*t*,1_) is negligible in comparison to *r*_*t*_ *- Q*(*s*_*x,t*_, *a*_*t*,2_) where *r*_*t*_ is either zero or one and *E*[*Q*(*s*_*x,t*_, *a*_*t*,2_)] = 0.5 and *Q*(*s*_*x,t*_, *a*_*t*,2_) > 0. So, if the (*α* + *λ*) is bigger than 0.4, which was the case for almost all of our subjects, the effect each individual trial has on |Δ*Q*| is bigger for the model-free agent. We computed the same |Δ*Q*| for other model-based and model-free formulations as well and this result held for a wide range of parameter values. In the next segment, we back up this result with simulation.

**Figure S1:**
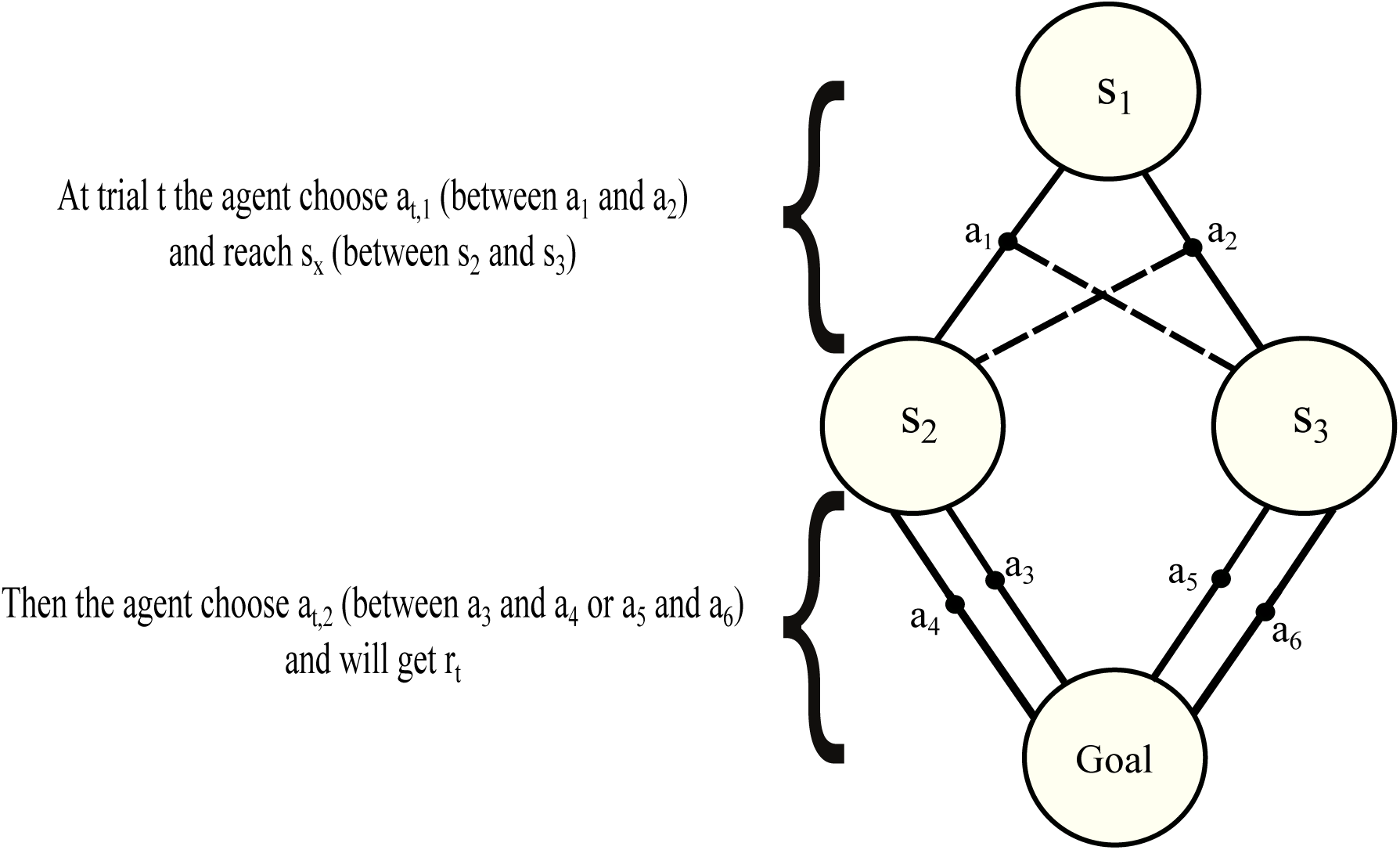
Diagram of two-step decision-making task

### Model-free strategy differentiate two first-step actions more extremely than model-based strategy

Model-based and model-free strategies have different methods to learn the values of first-step actions. To see how these two strategies differentiate between the two first-step actions, we simulated a pure model-based and a pure model-free agent, 100 times. We subtract the value of one action from the other to investigate the changes of this difference. The difference in action values is a confidence correlate. We did this analysis for different variabilities of the reward probabilities random walks. Confidence correlate for model-free strategy was significantly higher relative to model-based strategy for different variations of the reward generator (Figure S2). We checked this analysis for different values of *β* and *α* and the result was similar. Also, we checked this with other models for model-based and model-free learning and similar results were observed.

**Figure S2:**
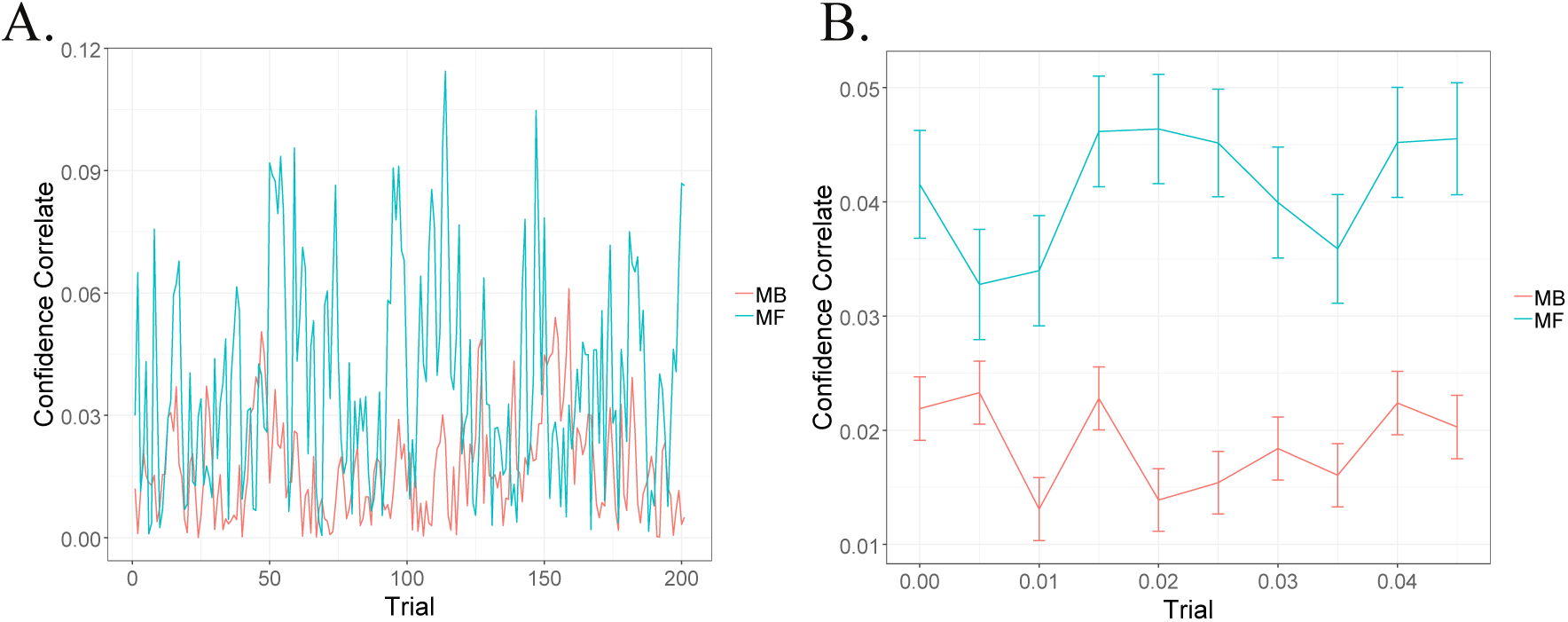
Confidence correlate of model-based and model-free strategies. A) confidence correlate for model-based (red) and model-free (green) strategies during trials. B) Mean confidence correlate of each agent during trials for different variabilities of reward generator.

